# Neural Representations of Visual Anagrams in Macaque Inferotemporal Cortex

**DOI:** 10.64898/2026.10.05.755559

**Authors:** Yipeng Li, Rui Zhang, Jingqiu Luo, Zishuo Zhao, Xieyi Liu, Zhengxin Gong, Wei Jin, Wanru Li, Pinglei Bao

**Affiliations:** School of Psychological and Cognitive Sciences, Peking University, Beijing 100871, China; Peking-Tsinghua Center for Life Sciences, Peking University, Beijing 100871, China; IDG/McGovern Institute for Brain Research, Peking University, Beijing 100871, China

**Keywords:** Inferotemporal cortex, Visual anagram, Object recognition, Transformation tolerance

## Abstract

Visual recognition requires stable representations of object identity despite large changes in the retinal image. Yet this stability must be selective: some transformations change not only how an image looks, but also what object it appears to depict. Visual anagrams highlight this challenge: a single image can resemble two different objects at different orientations, such as an elephant and a rabbit. Here, we asked how macaque inferotemporal (IT) cortex represents such transformations. We recorded large neuronal populations while presenting visual anagrams and unambiguous exemplars of their two interpretations across rotations. Neurons preferring one exemplar tended to prefer anagram orientations supporting the corresponding interpretation. At each of the two interpretation-relevant canonical angles, population responses more closely resembled those to the matching exemplar than those to the nonmatching exemplar. This matching advantage persisted when both exemplars were presented at the alternative canonical angle. Decoders trained exclusively on exemplars across rotations also tracked the shifts in anagram representations. Visual models showed anagram–exemplar alignment as well, with stronger alignment in models with greater relative shape preference. These findings suggest that transformation tolerance in IT is sensitive to what an image appears to depict, while relative shape preference is associated with similar representational organization in visual models.

## Introduction

Recognizing an object requires linking different retinal images to the same identity. Changes in position, distance, orientation, and illumination can substantially alter an object’s appearance, yet humans and other primates recognize objects rapidly across many such changes (Logothetis et al., 1994; Rajalingham et al., 2015, 2018). This capacity, commonly termed transformation tolerance, requires representations that preserve information about object identity across variations in visual input (DiCarlo et al., 2012; DiCarlo & Cox, 2007; Ito et al., 1995; Okamura et al., 2014). Understanding how the visual system achieves this tolerance is central to explaining how primates recognize the same object under the diverse viewing conditions encountered in natural vision.

Inferotemporal (IT) cortex plays a central role in supporting transformation-tolerant object representations. Along the ventral visual pathway, neuronal responses become increasingly selective for complex objects while showing greater tolerance to changes in position, scale, and viewpoint (Bao et al., 2020; Freiwald & Tsao, 2010; Kumar et al., 2019; Rust & DiCarlo, 2010). Whether such tolerance reflects abstract global-shape representations or combinations of diagnostic visual features remains debated (Ayzenberg & Behrmann, 2022). Regardless of the underlying mechanism, invariance cannot be applied indiscriminately. Too little invariance would fragment different views of the same object into separate identities, whereas too much invariance would collapse images that should be distinguished (Zoccolan et al., 2007). Successful recognition therefore requires the visual system to determine which image changes should be ignored and which indicate a genuine change in what object is depicted. Most previous studies have examined transformations of clear, unambiguous objects whose identity remains constant across the tested views (Bao et al., 2020; Chang & Tsao, 2017; Freiwald & Tsao, 2010; C. P. Hung et al., 2005; Kumar et al., 2019; Rust & DiCarlo, 2010). These studies reveal how IT generalizes across identity-preserving transformations but provide less insight into where this generalization should stop. In particular, it remains unclear how IT responds when a transformation that usually preserves identity instead changes what the image appears to depict.

Visual anagrams provide a distinctive test of this problem (Geng et al., 2024). In a rotational visual anagram, the same base image supports two alternative object interpretations at different orientations (Figure 1A). Rotation therefore changes not only the orientation of the image but also what object the image appears to depict. Behavioral studies in humans have shown that these orientation-dependent changes are reliable and can alter associated semantic judgments, including animacy and real-world size (Boger & Firestone, 2025, 2026). Unlike conventional rotation sequences, in which all views depict the same object, visual anagrams create a controlled conflict between transformation tolerance and sensitivity to changes in recognizable object interpretation: different views are related by a simple physical transformation, yet they are associated with different depicted objects. Visual anagrams therefore provide an opportunity to examine how high-level visual representations are reorganized when a normally identity-preserving transformation crosses an object-interpretation boundary.

**Figure 1.**
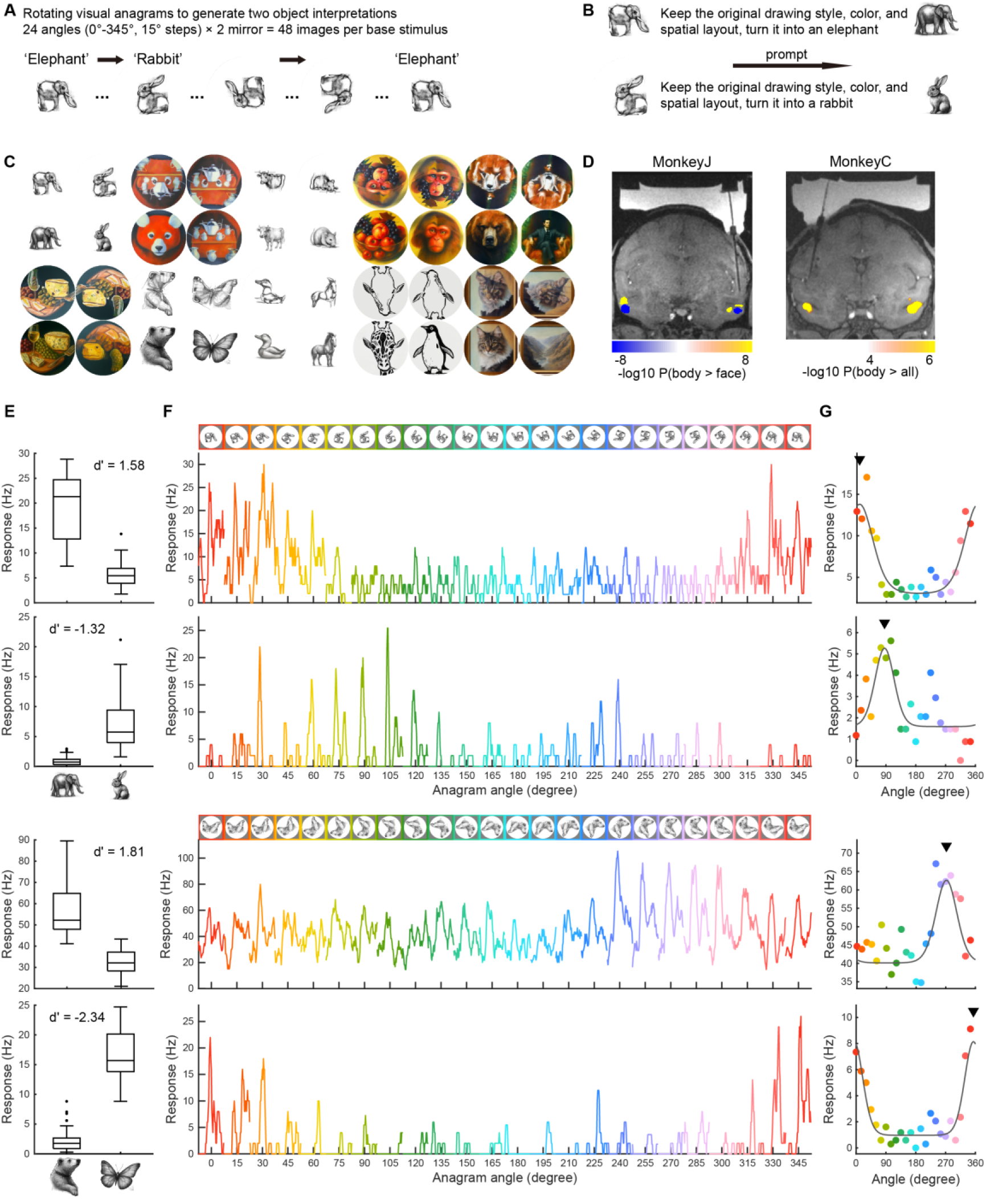
Stimuli, recording sites, and example single-unit tuning. (A) For each image, 24 rotation angles were generated and combined with two mirror conditions, yielding 48 variants per stimulus. (B) For each anagram, the image was rotated to the two canonical angles, and the resulting views were used to generate perceptually disambiguated exemplars. (C) All 10 anagram families. For each family, the anagram is shown at its two canonical angles, together with the corresponding canonical exemplars generated for the two interpretations. (D) Left, coronal slices showing fMRI-targeted anterior body (ASB) and face (AL) areas in Monkey J. Right, targeting of ASB in Monkey C. Activations for the contrast are shown as uncorrected P values on a signed log scale. (E–G) Four example exemplar-selective neurons. (E) Boxplots showing exemplar selectivity of the example units. (F) Neuronal tuning to 24 angles of anagrams. Top, anagram at 24 angles (colors indicate angles and were not visible to the monkeys during the experiment). Bottom, PSTHs of exemplar-selective neurons to anagrams, with each trace indicating firing rates from −50 to 450 ms after image onset. (G) Tuning curves of the example neurons. Black lines indicate von Mises fits to responses as a function of angle; triangles indicate the fitted preferred angles; colored dots indicate the angles shown in (F).

Angle-dependent responses alone, however, would not establish object-related reorganization, because rotating an image necessarily changes the retinal positions and orientations of its constituent visual features (Taubert et al., 2015). The key question is whether this response variation is related to independently measured object-related reference states. We therefore generated unambiguous canonical exemplars corresponding to the two recognizable interpretations of each anagram and measured neuronal responses to these exemplars (Figure 1B). Generic sensitivity to image orientation would not, by itself, predict a systematic relationship between anagram angle preferences and exemplar preferences measured across rotations. By contrast, if IT activity contained an object-related component, neurons preferring a given exemplar should tend to respond most strongly near its associated canonical angle, and population responses at that angle should more closely resemble the matching than the nonmatching exemplar state.

Here, we used visual anagrams to ask whether transformation tolerance in object vision is constrained by depicted identity. Macaques viewed anagrams and their corresponding canonical exemplars across densely sampled orientations while we recorded from fMRI-defined face- and body-selective regions of IT cortex. The canonical exemplars provided separately measured reference states for the two possible object interpretations of each anagram. Across single neurons, population geometry, and exemplar-trained decoders, rotation-dependent responses to anagrams were systematically organized with respect to these exemplar-defined states. We then extended the same framework to visual models as a computational probe of this organization. The model results showed that analogous anagram–exemplar alignment was strongest in models with greater relative shape-texture preference, linking this representational property to relative global shape preference in visual models.

## Results

### Neural tuning to visual anagrams aligns with preferred object interpretations

We selected 10 visual anagrams as base images, including 6 from examples provided in the anagram generation method paper (Geng et al., 2024) and 4 from a previous behavioral study (Boger & Firestone, 2025), with each anagram supporting two alternative object interpretations depending on its rotation angle (Figure 1A). To characterize neural preferences associated with each interpretation, we generated two perceptually disambiguated canonical exemplars for each anagram using a diffusion-based image generation model. For example, rabbit- and elephant-dominant views of the elephant-rabbit anagram were used to generate unambiguous rabbit and elephant exemplars, respectively (Figure 1B). We refer to these two interpretation-dominant orientations as the **canonical angles** of the anagram. This procedure yielded 20 exemplars corresponding to the two interpretations of the 10 anagrams; Figure 1C shows each anagram at its two canonical angles together with the corresponding canonical exemplars. Each image was presented at 24 angles spanning 0° to 345° in 15° increments, in random order. We additionally generated horizontally mirrored versions to reduce the contribution of contralateral visual-field preferences for local image features (Bao & Tsao, 2018; Issa & DiCarlo, 2012; Sharma et al., 2024). The resulting set comprised 30 base stimuli: 10 anagrams and 20 exemplars. Each was presented at 24 rotation angles in two mirror conditions, yielding 1,440 stimulus images. An additional set of 36 localizer images, comprising 12 faces, 12 bodies, and 12 objects, was used to characterize general category selectivity (Figure S1). Macaques were trained to maintain fixation on a dot presented on the screen to receive a juice reward, and stimuli were presented in random order for 300 ms each, followed by a 300 ms inter-stimulus interval. Neural activity was recorded using 45-mm Neuropixels NHP probes (Trautmann et al., 2025). In total, we obtained 13 recording sessions from 3 macaques. After applying unit inclusion criteria, the final dataset comprised 4,725 units with split-half reliability greater than 0.4 (Table S1). In the first 5 recording sessions, only 5 of the 10 anagram families were used in the experiment (see Methods).

In an initial experiment, we presented visual anagrams without corresponding canonical exemplars and recorded from face- and body-selective regions in middle and anterior IT (Figure S2A). For anagrams contrasting face- and body-related interpretations, units with different category preferences, defined using an independent localizer, favored different canonical angles. This relationship was more pronounced in anterior IT (Figure S2B), which also showed greater consistency of angle tuning across mirror conditions (Figure S2C-S2E, P < .001). These observations motivated us to focus the main experiment on the anterior lateral face patch (AL) and anterior STS body area (ASB; Figure 1D), and to use corresponding canonical exemplars to test how anagram angle preferences relate to preferences for the specific objects depicted.

We first asked whether a unit’s preference for the two canonical exemplars was associated with its angle tuning to the corresponding anagram. For each neuron and each anagram, we quantified canonical-exemplar selectivity as the *d’* between responses to the two corresponding canonical exemplars, each presented in 48 variants (24 rotation angles × 2 mirror conditions; Figure 1E). Units with |*d’*| > 0.5 were classified as canonical-exemplar-selective, with the sign of *d’* indicating the preferred exemplar.

Exemplar-selective units exhibited anagram angle preferences consistent with their exemplar preferences. For the elephant-rabbit anagram, for example, an elephant-selective unit responded most strongly near the elephant-associated canonical angle, whereas a rabbit-selective unit responded preferentially near the rabbit-associated canonical angle (Figure 1F, top). To quantify these preferences, we fitted each unit’s responses across the 24 anagram angles with a von Mises function (Swindale, 1998), which accounts for the circular nature of rotation angle (see Methods). The fitted mean direction (*μ*) defined the preferred angle. For these two example units, the estimated preferred angles were close to 0° and 90°, corresponding to the elephant- and rabbit-associated canonical angles, respectively (Figure 1G, top). A similar relationship between exemplar selectivity and anagram angle tuning was observed for the bear-butterfly anagram (Figure 1F, G, bottom). These examples illustrate angle preferences consistent with the object interpretation supported by each view and the neuron’s preference for the corresponding exemplar.

To determine whether the angle preferences observed in example units reflected a systematic population-level pattern, we next examined the distribution of preferred angles. Preferred angles for anagrams were strongly biased toward canonical angles at which the images supported recognizable object interpretations. For the elephant–rabbit anagram, many elephant-preferring units showed preferred angles near the elephant-associated orientation around 0° (mode = 7°; Figure 2A, 2B left), whereas rabbit-preferring units were biased toward the rabbit-associated orientation around 90° (mode = 95°; Figure 2A, 2C left). A similar pattern was observed for the bear–human anagram (Figure 2D–F, and other anagram examples in Figure S3), with modes at 178° for the bear-preferring population and 3° for the human-preferring population. To quantify this relationship across populations, for each qualified fit (see Methods), we calculated the circular distance between the preferred anagram angle and the canonical angle corresponding to its exemplar preference (“matching”, for example, distance between an elephant unit’s preferred angle and 0°), as well as the distance to the alternative canonical angle (“nonmatching”, for example, distance between an elephant unit’s preferred angle and 90°), and then averaged these distances within each population. Across populations, preferred angles were significantly closer to the matching than to the nonmatching interpretation-relevant canonical angles (Figure 2G; P < .001, Wilcoxon signed-rank test). These results indicate that, at the population level, IT angle preferences for visual anagrams are systematically organized around interpretation-relevant canonical angles defined by units’ preferences for the corresponding exemplars.

**Figure 2.**
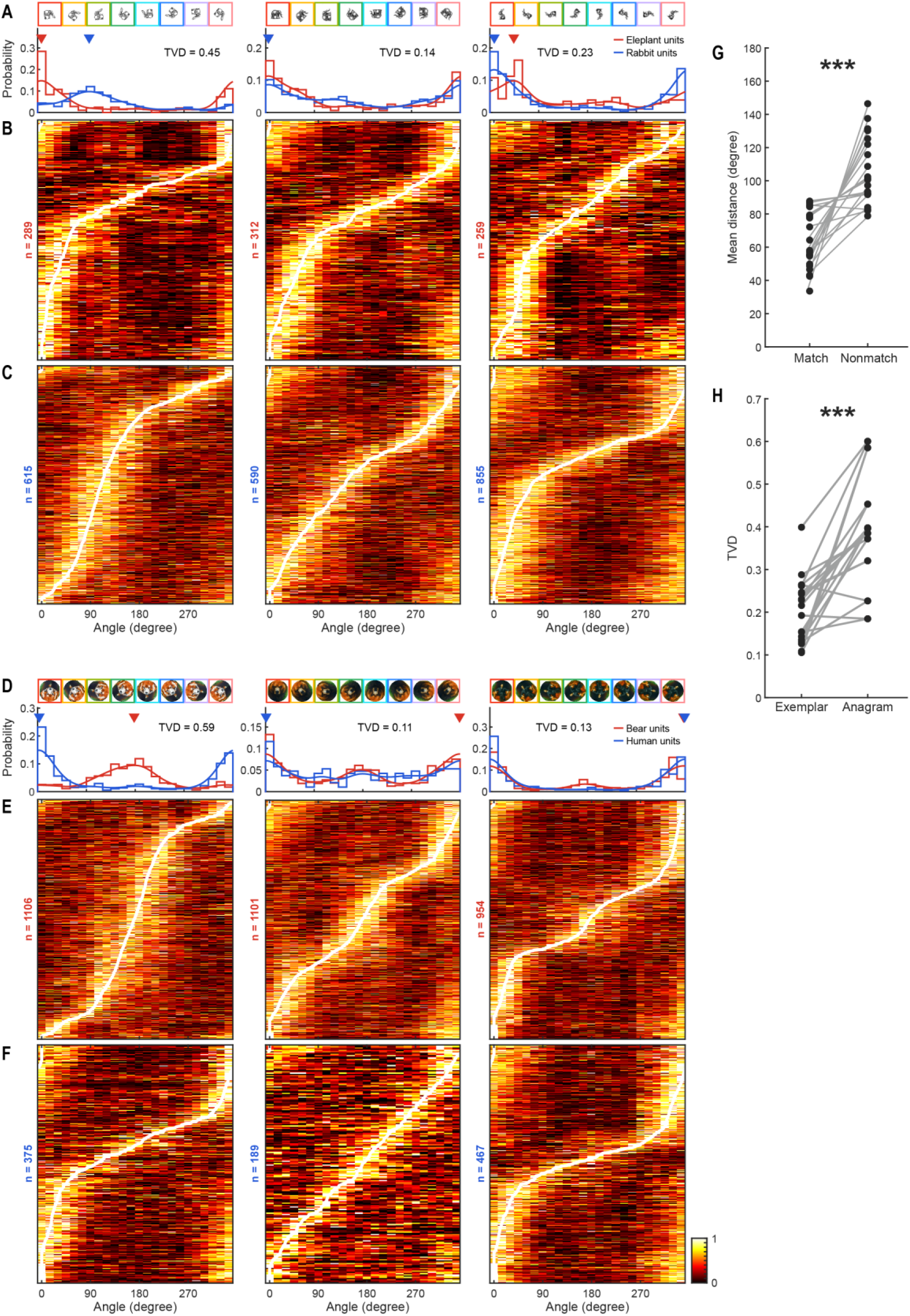
Rotation preferences of canonical-exemplar-selective neurons. (A–C) Example tuning summary for the elephant–rabbit anagram. From left to right, data are shown for the anagram, exemplar elephant, and exemplar rabbit. (A) Top, example images at eight rotation angles. Colored borders indicate angles. Bottom, distributions of fitted preferred angles for elephant-preferring neurons (red) and rabbit-preferring neurons (blue). Curves show kernel-density estimates, and triangles indicate the modes of the estimated distributions. TVD denotes the total-variation distance between two populations. (B) Heatmaps showing responses of the elephant preferring populations. Each row represents one qualified fit and shows its response across the 24 orientations, normalized between the minimum and maximum response within that row. Rows are ordered by fitted preferred angle. White lines indicate fitted preferred angles. (C) Same as (B) but for the rabbit preferring populations. (D–F) Same as (A–C), but for the bear–human anagram. (G) Mean circular distances of qualified fits from the matching and nonmatching canonical angles. Each connected pair represents one family-by-exemplar-preference population. (H) Separation between the preferred-angle distributions of the two exemplar-preferring populations, quantified using total-variation distance. *** indicates P < .001, Wilcoxon signed-rank tests

We next asked whether the separation between the two exemplar-defined subpopulations could be explained by a general difference in rotation preference. If so, the two subpopulations should also show distinct preferred-angle distributions when viewing rotated canonical exemplars. Instead, whereas the elephant- and rabbit-preferring subpopulations exhibited distinct preferred-angle distributions for the elephant–rabbit anagram, their distributions overlapped more strongly for both canonical exemplars, with both populations tending to prefer upright views. For example, rabbit-preferring units tended to have preferred angles near 0° when viewing the canonical elephant, where it was upright (Figure 2A–C, middle and right). We quantified the separation between the two preferred-angle distributions using total variation distance (TVD). For the elephant–rabbit family, TVD was 0.45 for the anagram, compared with 0.14 and 0.23 for the canonical elephant and rabbit exemplars, respectively. Across 20 family-by-exemplar comparisons drawn from the 10 anagram families, TVD was greater for anagrams than for the corresponding canonical exemplars (Figure 2H, P < .001, Wilcoxon signed-rank test). These results indicate that the two exemplar-defined subpopulations showed greater divergence in angle preference for anagrams than for their corresponding canonical exemplars. Thus, the separation observed for anagrams is unlikely to reflect only a general difference in rotation preference between the two subpopulations.

### Population representations of visual anagrams align with corresponding object exemplars

The preceding analyses identified exemplar-aligned angle tuning within neuronal subpopulations defined by their responses to the canonical exemplars. We next asked whether this organization was also evident in the entire recorded population, without selecting units based on exemplar preference. For each anagram, we constructed a population response vector at each of the 24 rotation angles. Responses to the two horizontal-mirror conditions were concatenated as separate feature dimensions, yielding a 2N-dimensional vector in which both mirror configurations contributed equally.

We first computed an angle-by-angle representational dissimilarity matrix (RDM) for each anagram using Spearman correlation distance. The RDMs were broadly banded, consistent with population representations that changed smoothly with rotation angle, but also showed localized departures from this organization at particular angles (Figure 3A). To determine whether this angle-dependent geometry was organized relative to two corresponding exemplar states, we expanded the analysis to include the 24 angle-specific responses to each of the two corresponding canonical exemplars, yielding 72 conditions in total. We then visualized the resulting geometry using multidimensional scaling (MDS). In these visualizations, the anagram trajectory approached different canonical-exemplar trajectories near the two corresponding canonical angles (Figure 3B; additional examples in Figure S4). For example, responses to the red panda–kitchen table anagram near 0° were positioned closer to the red panda exemplar trajectory, whereas responses near 180° were positioned closer to the kitchen table exemplar trajectory.

**Figure 3.**
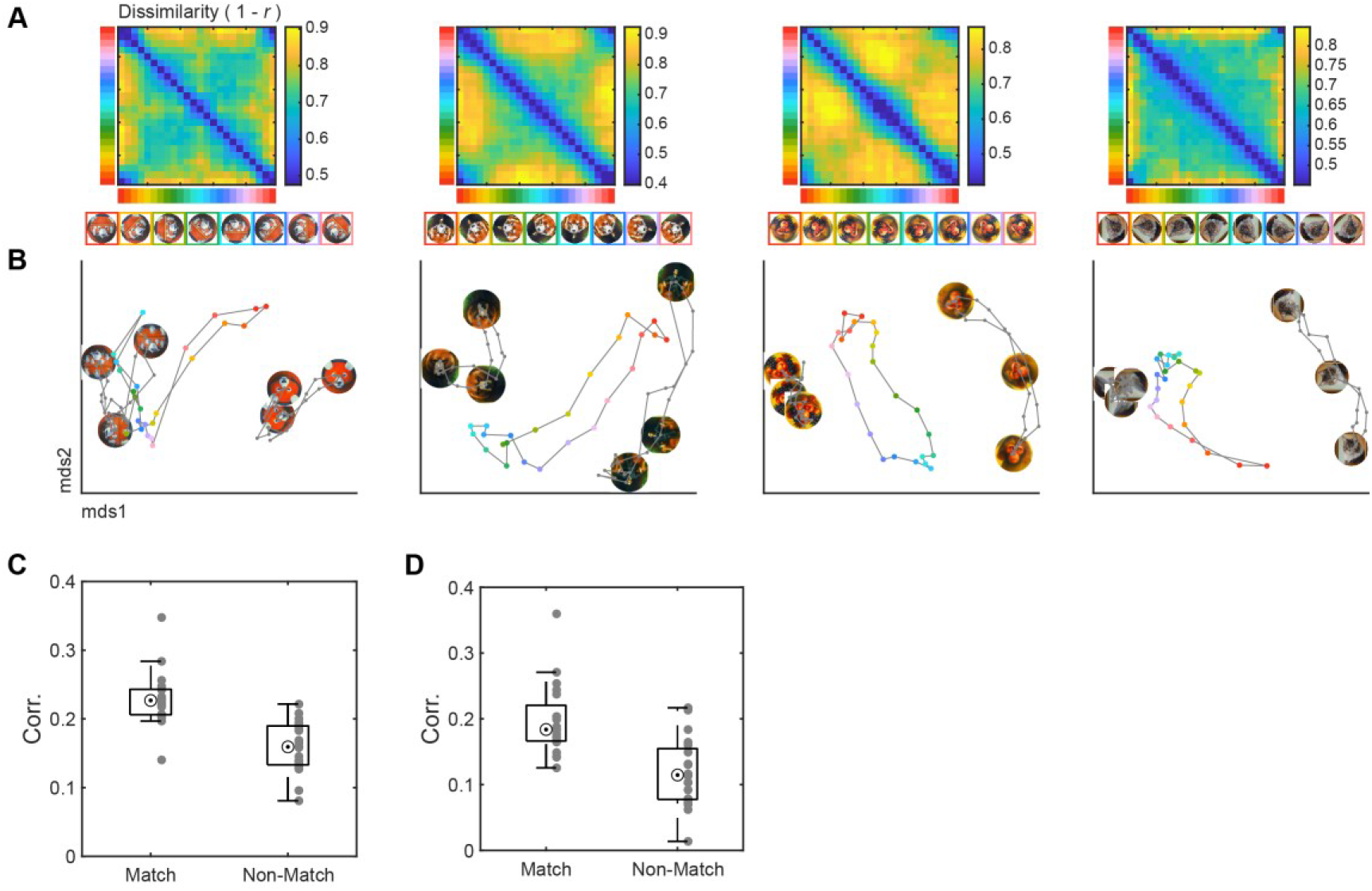
Population representational geometry across anagram rotations. (A) Top, RDMs of IT population responses to four example anagrams presented at 24 rotation angles. Dissimilarity was measured as Spearman correlation distances. Colored bars along the axes indicate rotation angle. For display, diagonal value was set to the minimum off-diagonal dissimilarity. Bottom, anagram stimuli at selected angles. (B) MDS visualization of the population geometry for each anagram family. Each analysis included responses to the anagram and its two corresponding canonical exemplars, each presented at 24 rotation angles, yielding 72 conditions. Colored dots and lines trace the anagram response trajectory across rotation angles, whereas gray trajectories show responses to the two exemplars. Enlarged stimulus images mark selected positions along the exemplar trajectories. (C) Mean correlations between population responses to anagrams at their canonical angles and responses to the matching or nonmatching exemplar across rotations. For each interpretation-specific comparison, the population response vector for the anagram at its canonical angle was correlated separately with the response vector for each exemplar at each of the 24 rotation angles. The resulting 24 correlations were then averaged for each exemplar. Each gray dot represents one interpretation-specific comparison. (D) Correlations between each canonical-angle anagram response and responses to the matching and nonmatching exemplars, with both exemplars presented at the alternative canonical angle associated with the nonmatching interpretation. Each gray dot represents one interpretation-specific comparison.

Because MDS provides only a low-dimensional visualization, we quantified this alignment directly in the original population space. For each of the 20 interpretation-specific conditions (two per anagram), we constructed a population response vector for the anagram at the corresponding canonical angle. We then calculated its correlation separately with the population response to each of the two exemplars at each of the 24 rotation angles and averaged the resulting correlations across angles. The mean correlation was significantly higher for the matching exemplar than for the nonmatching exemplar (Figure 3C; t(19) = 4.59, P < .001).

We next asked whether the matching advantage persisted when the matching exemplar was rotated away from its canonical orientation. For each anagram at a canonical interpretation-relevant angle, we compared its population response with responses to a rotated matching exemplar and an unrotated nonmatching exemplar. For example, for the bear–human anagram presented at its bear-associated canonical angle, we correlated its population response vector with those of the bear exemplar (matching) and the human exemplar (nonmatching), both presented at the human-associated canonical angle (note that, in this condition, the bear exemplar was rotated by 180° from its upright orientation and was therefore inverted, while the human exemplar remained upright at 0°). The correlation remained significantly higher for the matching exemplar (Figure 3D; t(19) = 3.78, P < .001), showing that the matching advantage persisted when the anagram and exemplars were presented at different rotation angles. These analyses show that angle-dependent variation in IT population activity is systematically aligned with canonical-exemplar states associated with the interpretation supported by each view, even when the matching exemplar was rotated away from its upright orientation and the nonmatching exemplar remained upright, indicating that this alignment did not require identical image orientations.

### Population decoding reveals angle-dependent shifts along a canonical-exemplar axis

MDS visualizations suggested that anagram responses approached different canonical-exemplar states across rotation angles, but did not directly quantify their relative correspondence to the two exemplars. We therefore trained a ridge-regularized logistic decoder within each recording session to discriminate population responses evoked by the two canonical exemplars and applied the decoder to responses evoked by the corresponding anagram from the same session (Figure 4A). Critically, the decoder was trained exclusively on responses to the canonical exemplars, with each class represented by the complete set of 48 variants (24 rotation angles × 2 horizontal-mirror conditions). Thus, the decoder should learn the differences between the two exemplars that generalized across the sampled rotations and mirror configurations, without using anagram responses during decoder training. Decoder output was expressed as the probability assigned to exemplar A: values approaching 1 indicated stronger evidence for exemplar A, values approaching 0 indicated stronger evidence for exemplar B, and values near 0.5 indicated approximately equal evidence for the two classes.

**Figure 4.**
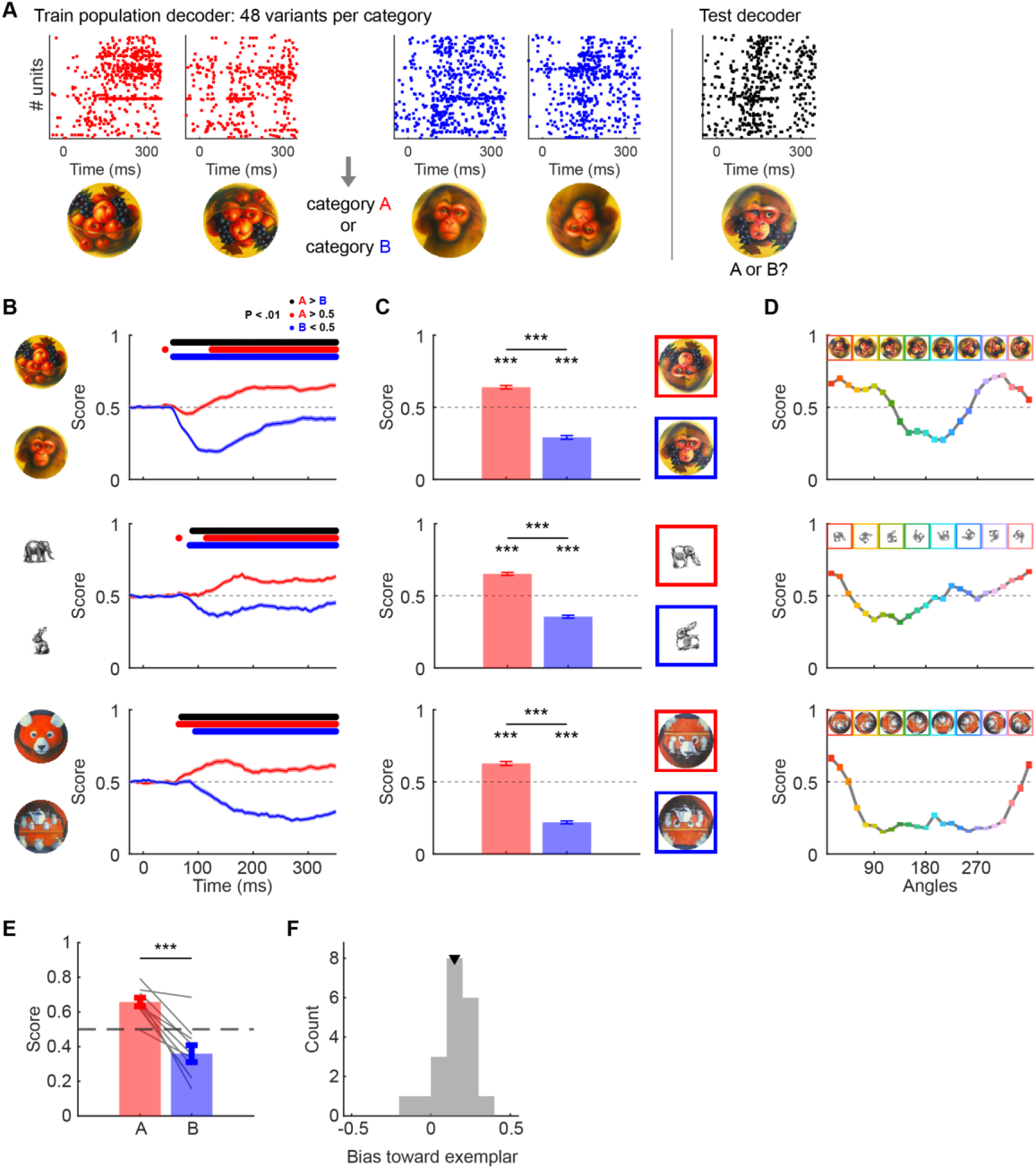
Population decoding for visual anagrams. (A) Decoding approach. Population responses were used to train a logistic decoder to discriminate between two exemplars. The trained decoder was then applied to responses evoked by the corresponding visual anagram. In (B–E), decoder scores greater than 0.5 indicate alignment with the exemplar assigned as 1 during training, whereas values below 0.5 indicate alignment with the exemplar assigned as 0. (B) Time-resolved decoder scores for the fruit bowl-monkey face anagram (top), elephant–rabbit anagram (middle), and red panda–kitchen table anagram (bottom). Red and blue traces show responses averaged across three neighboring angles centered on the canonical angles associated with exemplars A (fruit bowl, elephant and red panda) and B (monkey face, rabbit and kitchen table), respectively. The top black bar indicates time points at which decoding score for the red angle is significantly greater than for the blue angle; the red bar indicates time points at which the red trace is significantly greater than 0.5; the blue bar indicates time points at which the blue trace is significantly less than 0.5 (FDR-corrected P < .01). (C) Time-averaged (60-400 ms) decoding score for the same example pairs shown in (B). (D) Decoding score as a function of angle for the three example anagram pairs. (E) Decoding scores across the 10 anagram pairs. Each line represents one anagram pair. Higher values indicate decoding score for the ‘1’ condition relative to the ‘0’ condition during exemplar training, “A” and “B” correspond to anagrams shown at the canonical angles associated with exemplar A or exemplar B, respectively. (F) Distribution of decoder bias toward the exemplar matching each canonical angle across 20 interpretation-specific conditions. Bias was calculated as score − 0.5 for A-associated canonical angles and 0.5 − score for B-associated canonical angles, such that positive values indicate greater evidence for the matching exemplar. Shaded regions in (B) and error bars in (C-D) indicate the SEM across trials; error bars in (E) indicate the SEM across anagram families. Markers in (F) denote mean values. Scores in panels B, C, E, and F were pooled across the three angles centered on each canonical angle. *** indicates P < .001.

Decoder evidence varied systematically with anagram angle. For the fruit bowl-monkey face anagram, 3 angles near the fruit bowl-associated canonical angles (345°, 0°, and 15°) produced stronger evidence for the fruit bowl exemplar class, whereas angles near the monkey face-associated canonical angles (165°, 180°, and 195°) shifted decoder output toward the monkey face exemplar class (Figure 4B–D; additional examples in Figure S5). Across the 10 anagram families, decoder scores shifted in the expected direction between neighborhoods centered on the two canonical angles (Figure 4E; t(9) = 6.74, P < .001). When decoder output around each canonical angle was recoded as evidence for the matching exemplar, it was also significantly biased above 0.5 (Figure 4F; t(19) = 5.57, P < .001). The magnitude of this modulation varied across stimuli: the duck–horse and table–turtle anagrams, for example, were dominated by the duck or turtle exemplar class across most rotation angles (Figure S5), potentially reflecting differences in stimulus quality or the selectivity of the recorded neurons.

Overall, these results show that angle-dependent anagram responses vary systematically along a population axis defined by the two canonical exemplars, in a direction consistent with the recognizable interpretation associated with each view.

### Anagram–exemplar alignment in visual models is associated with relative shape preference

The exemplar-trained decoders revealed systematic rotation-dependent shifts in IT responses to visual anagrams. We next asked whether visual models exhibited analogous anagram–exemplar alignment, using methods similar to Figure 4. We applied the same decoding framework to representations from 37 models: for each model and anagram family, a decoder trained to distinguish the two canonical exemplars across rotations and mirror conditions was applied to representations of the corresponding anagram. For each canonical angle, decoding bias was defined relative to 0.5 and oriented such that positive values indicated greater evidence for the matching exemplar. Biases were then averaged across the two canonical angles and anagram families to obtain a matching-exemplar decoding bias for each model. Across models, the mean bias was 0.19. For descriptive comparison, applying the corresponding analysis to a neural pseudo-population constructed by pooling units across sessions yielded a mean bias of 0.26 (Figure 5A).

**Figure 5.**
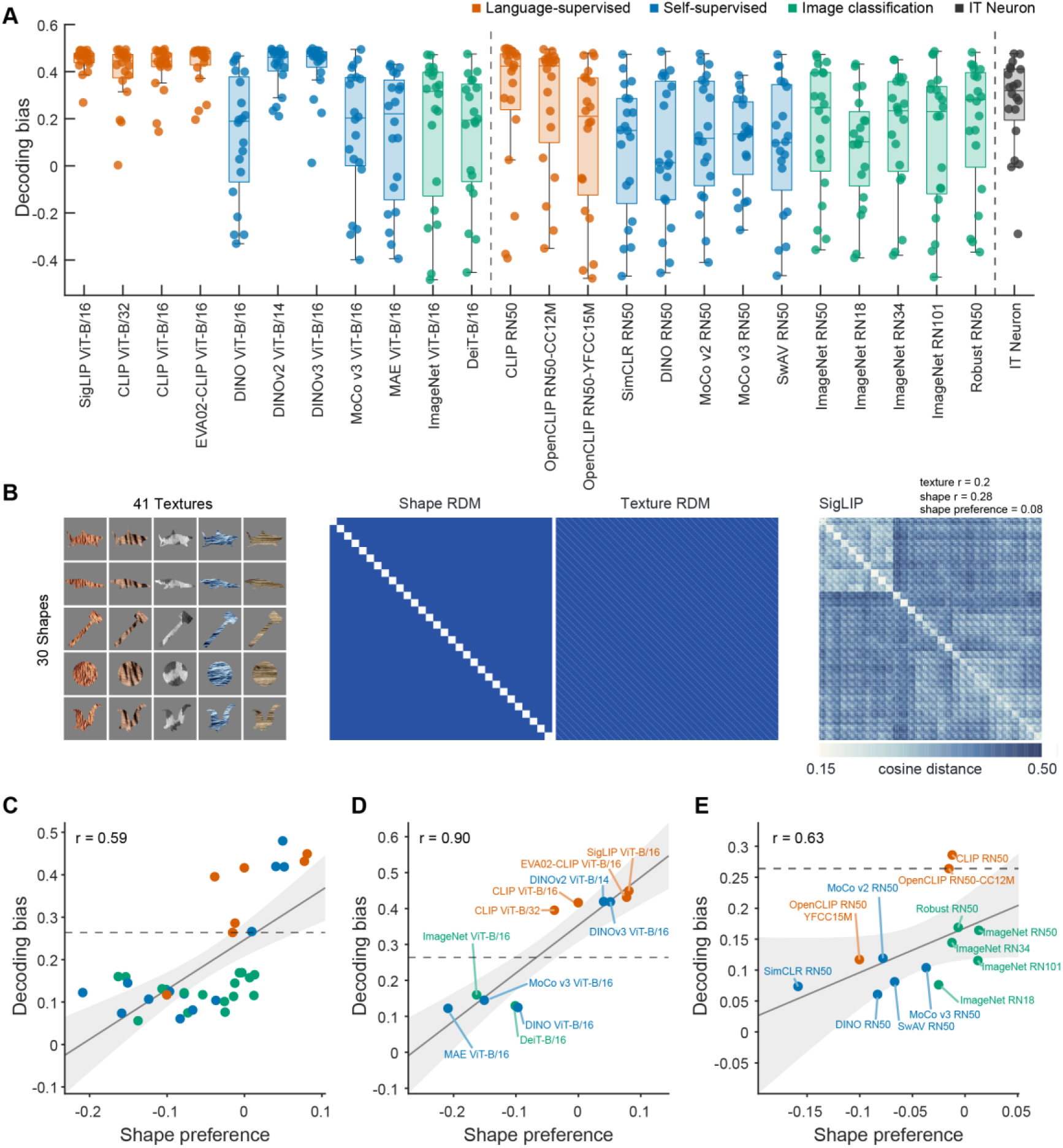
Canonical-exemplar decoding of visual anagrams across visual models. (A) Matching-exemplar decoding bias for 11 ViT-B models, 13 ResNet models, and the IT neuronal population. Decoders were trained to distinguish the two canonical exemplars and then applied to anagrams at their two interpretation-relevant canonical angles. Each point represents one of the 20 interpretations. (B) Procedure for quantifying relative shape versus texture preference. Left, example stimuli illustrating combinations of shapes and textures. Middle, theoretical representational dissimilarity matrices (RDMs) for shape and texture. Right, example model RDM together with its corresponding partial correlations with the shape and texture RDMs. (C) Relationship between relative shape preference and matching-exemplar decoding bias across all 37 models. Orange denotes language-supervised models, blue denotes self-supervised models, and green denotes image-classification models. (D) Same relationship for the 11 ViT-B models. (E) Same relationship for the 13 ResNet models. The neuronal value, derived from pooled data across recording sessions, is shown in gray in panel A and as a horizontal reference line in panels C–E. Each point in C–E represents one model.

Matching-exemplar bias varied substantially across models, motivating us to ask whether this variation was related to models’ relative reliance on shape versus texture. Previous work has shown that standard ImageNet-trained CNNs often rely more strongly on texture than human observers and that relative shape–texture preference varies with architecture, training objective, and training procedure (Baker et al., 2018; Geirhos et al., 2022; Hermann et al., 2020). This distinction is particularly relevant for visual anagrams, because the same image content supports different recognizable object interpretations at different orientations. Therefore, we quantified each model’s relative shape preference using 1,230 images that factorially combined 30 shapes with 41 textures (Figure 5B; see Methods). For each model, we calculated the partial Spearman correlation between its representational dissimilarity matrix and a shape-based hypothesis matrix while controlling for texture, as well as the complementary partial correlation with texture while controlling for shape. Relative shape preference was defined as the partial shape correlation minus the partial texture correlation, with positive values indicating stronger shape-related than texture-related representational structure. Across all 37 models, relative shape preference was positively associated with matching-exemplar decoding bias (r = 0.59, P < .001; Figure 5C). This association was also observed within Vision Transformer Base models (ViT-B, r = 0.90, P < .001; Figure 5D) and within ResNet models (r = 0.63, P < .05; Figure 5E), showing that this relationship was present within both broad architectural families.

Because decoding measures information available to a linear readout, we also asked whether a corresponding relationship was evident at the level of individual model units, using methods similar to Figure 2. Among units selective for one canonical exemplar, we measured the circular distance between the preferred anagram angle and the canonical angles associated with the matching and nonmatching exemplars. We defined a normalized matching-angle advantage as the difference between the nonmatching and matching distances divided by the circular separation between the two canonical angles. Positive values therefore indicated that a unit preferred an anagram angle closer to the canonical angle associated with its preferred exemplar. IT neurons showed a positive normalized matching-angle advantage (mean = 0.30; Figure S6A). Across models, the mean unit-level matching-angle advantage was positively associated with relative shape preference (r = 0.50, P < .01; Figure S6B). This relationship was particularly strong within ViT-B (r = 0.86, P < .01; Figure S6C), whereas the positive association within ResNets did not reach statistical significance (r = 0.43, P = .15; Figure S6D).

Across the tested models, stronger anagram–exemplar alignment was associated with greater relative shape preference. This relationship was evident in population decoding and in individual-unit angle tuning. These analyses establish an association between the two representational properties but do not show that relative shape preference causes anagram–exemplar alignment or that the neural and model effects arise from the same computational mechanism.

The effects we reported here were generally consistent in direction across animals. In each of the three monkeys, preferred angles were significantly closer to the matching than the nonmatching interpretation (Figure S7A). The greater separation of preferred-angle distributions for anagrams than canonical exemplars was significant in Monkeys J and C and showed the same numerical direction but was not significant in Monkey Z (Figure S7B), for which fewer comparisons passed the fit-selection criteria.

Population responses were more strongly correlated with the matching exemplar representation (Figure S7C), and decoding was significantly biased toward the expected interpretation (Figure S7D). Thus, the principal interpretation-aligned tuning, representational-geometry, and decoding effects were not driven by a single animal.

## Discussion

The present results show that rotation-dependent responses in macaque IT are systematically organized relative to independently measured object-exemplar states. Across exemplar-selective neurons, population geometry, and exemplar-trained decoding, anagram orientations associated with a particular interpretation elicited activity more closely aligned with the corresponding canonical exemplar. This alignment persisted when anagrams and exemplars were presented at different angles and was stronger for anagrams than expected from the general rotation preferences of the canonical objects. Visual models showed an analogous representational signature, with stronger alignment in models exhibiting greater relative shape preference. Together, these findings suggest that IT does not treat transformation-related image variation independently of what the resulting image can depict.

These findings extend an emerging literature using visual anagrams to dissociate image content from recognizable object interpretation. Visual anagrams were introduced as images that support different recognizable interpretations when subjected to transformations such as rotation or reflection (Geng et al., 2024). Subsequent human studies showed that these orientation-dependent interpretations carry high-level object properties into behavior. Real-world size influenced familiar-size interference and preferred viewing size of the anagram (Boger & Firestone, 2025), while animacy influenced visual working memory and visual search (Boger & Firestone, 2026). The present study extends this behavioral work by showing that anagram rotations are accompanied by systematic changes in macaque IT cortex activity relative to independently measured exemplar states. The neural results therefore provide a representational counterpart to the behavioral observation that rotating an anagram can change more than its retinal orientation.

Visual anagrams also connect transformation tolerance to previous work on perceptual ambiguity. During binocular rivalry, activity in macaque IT can follow the animal’s reported or inferred percept (Hesse & Tsao, 2020; Sheinberg & Logothetis, 1997). Human studies have similarly shown that activity in category-selective ventral temporal cortex tracks the dominant interpretation of rivalrous face and house images (Tong et al., 1998, 2006) and the face versus vase interpretation of the Rubin figure (Andrews et al., 2002; Hasson et al., 2001). Another study used ambiguous photographs generated by morphing between two objects to show that responses of IT neurons to identical ambiguous images varied with the monkeys’ behavioral interpretations (Liu & Jagadeesh, 2008). The current study addresses a related but distinct question. In those studies, neural activity was related to trial-by-trial reported or inferred perception while the physical input was held constant or closely matched. Here, rotation changed the retinal image, and the macaques did not report their percepts. The canonical exemplars instead provided independently measured reference states for asking how a continuous transformation trajectory was organized when different portions of that trajectory supported different recognizable objects. Visual anagrams therefore bridge studies of transformation tolerance and perceptual ambiguity, while stopping short of demonstrating percept-dependent neural activity.

An important question is what type of visual information produces this object-related representation. Classic work characterized IT neurons as selective for moderately complex diagnostic features and proposed that objects are represented through combinations of feature-selective neuronal populations (Tanaka, 1996; Tsunoda et al., 2001). Recent evidence further suggests that feature selectivity can be spatially structured, with different facial features eliciting maximal responses at distinct receptive fields (Livingstone et al., 2026). Thus, rotating an anagram might bring diagnostic features, such as eyes, into the preferred spatial locations of neurons responsive to the corresponding canonical exemplar. However, IT neurons can also integrate multiple contours and encode complex shape configurations (Brincat & Connor, 2004, 2006; C.-C. Hung et al., 2012; Yamane et al., 2008). Even facial identity can be represented through continuous visual-feature dimensions or populations driven by local facial features (Chang & Tsao, 2017; Waidmann et al., 2022). Such interactions between feature identity and spatial position could contribute to the observed interpretation-aligned tuning without requiring an explicit representation of perceived object identity. The present findings therefore do not resolve the distinction between visual-feature and object-identity accounts. The persistence of matching-exemplar alignment across different angles argues against a generic preference for absolute orientation, but it does not establish an identity code independent of visual form. Instead, a parsimonious interpretation is that diagnostic features and their spatial configurations may bias IT population activity toward states associated with the corresponding canonical exemplars. Object-related organization may therefore emerge from visual features whose diagnostic value depends on their orientation and global configuration.

The model analysis extends this account by identifying relative shape preference as a representational property associated with object-consistent organization. Across 37 models, matching-exemplar alignment was stronger in models whose representations emphasized shape over texture in an independent stimulus set. This relationship remained evident within both ViT-B models and ResNet models, showing that it was not driven simply by the distinction between transformer and convolutional architectures. The same organization was also echoed at the level of individual model units’ preference. These findings connect anagram–exemplar alignment to evidence that artificial visual systems differ substantially in their reliance on local texture versus global shape (Baker et al., 2018; Geirhos et al., 2022; Hermann et al., 2020). They also complement the approach used by another study (Doshi et al., 2025), which used patch-rearranged anagrams to demonstrate differences among models in configural-shape preference. Whereas that work examined whether models could recognize objects constructed from different arrangements of the same local elements, our analysis shows how rotating a single image shifts model representations toward independently defined exemplar patterns, and how this alignment relates to shape preference. Several models with the strongest alignment were language-supervised, further raising the testable possibility that language-aligned training promotes more object-structured visual representations. More broadly, these analyses move beyond asking which model best matches neural responses and identify relative sensitivity to shape as a representational property that helps explain why some visual systems organize transformations in an object-consistent manner.

Several limitations define the scope of the conclusions. First, because the macaques passively fixated, we cannot conclude that they consciously recognized the intended objects or experienced orientation-dependent perceptual change. The graded decoder outputs and continuous representational trajectories also do not establish subjective categorical boundaries. Demonstrating percept-dependent coding would require measuring macaques’ object judgments at each angle and testing whether neural alignment predicts their behavioral choices. Another consideration is our recording locations: our recordings were from anterior IT, whereas more anterior and medial regions show greater visual invariance and may represent later stages of object coding (Bao et al., 2020; Freiwald & Tsao, 2010). Downstream regions such as MTL and PFC may further stabilize and interpret visual representations in a more categorical manner (Cao et al., 2026; Freedman et al., 2001). Future recordings from these areas could test whether interpretation-specific states become more stable and more directly linked to perception and behavior.

In summary, visual anagrams reveal a dimension of transformation tolerance that is largely absent from studies of unambiguous objects. IT preserves information about how an image can change while organizing that variation relative to the different objects the transformed image can depict. This organization is consistent with sensitivity to diagnostic and configural shape information, is reflected to varying degrees in artificial visual models, and may depend on the visual experience that shapes which object properties are preserved or inferred from sensory input (N. Li & DiCarlo, 2010; Zhao et al., 2024). Visual anagrams therefore provide a means to investigate not only how invariant object representations are constructed, but also how the visual system preserves the distinctions that invariance should not erase.

## Data and code availability

Preprocessed neural responses and analysis code will be made publicly available upon publication.

## Methods

### Animals

All experimental protocols followed the guidelines of the Institutional Animal Care and Use Committee (IACUC) of the Peking University Laboratory Animal Center and were approved by the Peking University Animal Care and Use Committee (id: Psych-BaoPL-1). Six male rhesus macaques (Macaca mulatta), aged 6–8 years and weighing 6.5–11 kg, were used in this study. Three macaques (J, Z, and C) contributed to the main experiments, while five macaques (J, Z, F, L, and M) participated in the initial experiment shown in Figure S2. Monkey training and details of the implant or craniotomy surgery followed previously reported protocols (Y. Li et al., 2026) except that monkeys F and L were implanted with larger recording chambers. A summary of the recording locations is provided in Table S1.

### Stimuli generation and behavioral control

In the initial experiment shown in Figure S2, we used 12 visual anagrams as base images, of which 4 were taken from a previous behavioral study (Boger & Firestone, 2025) and 8 were from the examples provided with the original visual anagram paper (Geng et al., 2024). In the main experiment, we selected 10 of them to generate two canonical versions of each using a diffusion-based approach. For example, we rotated the ‘Monkey-Fruit’ anagram to “Monkey” angles and applied the prompt: “Keep the original drawing style, spatial layout, and colors, turn it into a monkey face.” and rotated the images to “fruit bowl” angles and applied the prompt: “Keep the original drawing style, spatial layout, and colors, turn it into a fruit bowl.” This procedure generated three base images from each original anagram, yielding 30 base images in total. For each stimulus, we generated 24 angles spanning 0–345° in 15° increments and applied a horizontal flip operation, resulting in 48 variants per base stimulus. Additionally, we used 12 face images, 12 body images, and 12 object images as localizer stimuli to identify unit selectivity (Figure S1). In total, the main experiment comprised 1,476 stimuli. During the first five sessions, only half of the stimulus set was presented (5 of 10 stimulus pairs; 756 stimuli in total). In the remaining sessions, the full set of 1,476 stimuli was presented (see Table S1 for details). Note that, since each anagram corresponds to two distinct canonical exemplars, we used 20 exemplar-specific comparisons derived from 10 anagram families as the statistical units in Figures 2G–H, 3C–D, and 4F.

Stimulus presentation, eye movement control, and reward delivery were controlled using the NIMH MonkeyLogic (Hwang et al., 2019) toolbox. Stimuli were shown on an LCD screen. Eye movements were monitored with infrared eye trackers (JSMZ). The macaques were trained to maintain fixation on a small fixation dot to receive juice reward. Stimuli were presented once the macaque had maintained fixation for at least 300 ms. In the initial experiment, images were presented with a 500 ms on / 250 ms off paradigm. In the main experiment, a 300 ms on / 300 ms off paradigm was used. Stimuli were shown in randomized order at a visual angle of 10°.

### fMRI-guided high-density electrophysiological recordings

We used Neuropixels NHP (Trautmann et al., 2025) probes to record from category-selective regions of IT cortex. Recording sites were guided by fMRI mapping of the monkey while viewing blocks of categorical images. The probe was advanced through a stainless-steel guide tube to facilitate dura penetration with a Narishige hydraulic micromanipulator. Probe advancement speed was reduced once signal silence in the STS transitioned into spiking activity, serving as a marker for entry into IT cortex. Neural activity was recorded using SpikeGLX at approximately 30,000 Hz. Data preprocessing was performed using SpikeInterface (Buccino et al., 2020), including 300 Hz high-pass filtering, removal of bad channels, sampling phase correction, and global median common referencing. Probe drift was corrected using DREDge (Windolf et al., 2025). Spike sorting was carried out with Kilosort4 (Pachitariu et al., 2024). Sorted units were automatically curated using BombCell (Fabre et al., 2023), and spike times were aligned to stimulus onset as measured by a photodiode.

We further included units that satisfied all of the following criteria: (1) a significantly higher firing rate during the 60–300 ms post-stimulus period compared to the baseline window (-25 to 25 ms), assessed using a Wilcoxon rank-sum test (P < .05); (2) a presence ratio larger than 0.95; (3) a waveform amplitude exceeding 20 μV; and (4) split-trial-half reliability above 0.4 after Spearman-Brown correction across the full image set (Van Bree et al., 2025). The resulting yields are reported in Table S1. For analyses of exemplar, face, or body selective units, a d-prime threshold of 0.5 was applied.

To facilitate interpretation, anagrams for which data were collected from the left hemisphere were left– right flipped during analysis and visualization, ensuring that stimulus features were aligned across hemispheres. Canonical angles were defined as the angles at which each anagram aligned with one of the two specific interpretations during stimulus generation.

### Single-unit tuning

In the main experiment, responses were quantified as spike counts measured from 60–400 ms after stimulus onset. Units selective for one of the alternative interpretations of anagrams were identified by computing d′ between the two corresponding canonical exemplars, based on responses to the 48 variants of each base stimulus. Units with |d′| > 0.5 were classified as exemplar-selective. The statistical significance of rotation or mirror tuning was evaluated using a one-way ANOVA across the 48 stimulus variants belonging to the same base stimulus. The resulting P-values were corrected across neurons using the FDR correction.

For each neuron and mirror condition, responses across the 24 rotation angles were fitted separately with a von Mises function. The model contained four parameters: the preferred angle, denoted as *μ*; a concentration parameter; a response-scaling parameter; and a baseline-response parameter. Parameters were estimated by bounded nonlinear least squares using MATLAB’s lsqcurvefit. After fitting, the model was evaluated at the 24 angles. Fit quality was defined as the correlation between the predicted tuning curve and the observed mean response across the 24 angles. The fitted *μ* parameter was used as the neuron’s preferred angle. For the preferred-angle analysis in Figure 2, a mirror-specific tuning fit was retained when tuning was significant after FDR correction (P < .001), and the correlation between the fitted and observed tuning curves was at least 0.3.

Each anagram had two defined interpretation-relevant canonical angles associated with the two canonical exemplars. For every qualified fit defined above, the shortest circular distance was calculated between the fitted *μ* and each of the two canonical angles. For example, for an exemplar A-selective neuron, the distance to the exemplar A canonical angle was defined as the matching distance, and the distance to the exemplar B canonical angle was defined as the nonmatching distance. Alignment was summarized using the mean matching and nonmatching distances of 20 exemplar-preferring populations and compared using a Wilcoxon signed-rank test.

Preferred-angle distributions were estimated separately for the two populations and for anagrams and canonical exemplars. Each qualified unit contributed a circular von Mises kernel centered on its fitted preferred angle; kernels were averaged within each population and normalized to unit integral. Separation between the two distributions was quantified using total-variation distance (TVD), with larger values indicating less overlap. TVD values were compared between anagrams and canonical exemplars using Wilcoxon signed-rank test.

In the initial experiment shown in Figure S2, responses to anagrams were quantified from 60–550 ms after stimulus onset. For the preferred-angle distributions in Figure S2B, each mirror orientation was fitted separately. A fit was included when the one-way ANOVA across all 48 variants of its anagram yielded an FDR-corrected P < .001, its von Mises fit correlation was at least 0.3, its face or body selectivity was at least 0.5. In Figure S2C-S2E, to assess consistency between mirror orientations, a unit was included when the FDR-corrected 48-variant ANOVA yielded P < .001 and both mirror-specific fits had correlations of at least 0.3. For each of the 12 anagrams, preferred angles from the two fits were compared across units using a circular–circular correlation. The correlations were then compared between middle and anterior IT across anagrams using Wilcoxon signed-rank test.

### Population analyses

RDM and MDS analysis: in Figure 3, neural responses were computed within a 60–400 ms window after stimulus onset for each stimulus. Trial-averaged neuronal responses from all sessions were concatenated into a pseudo-population. Within each anagram family, each of the three stimulus sets: anagrams, exemplar A, and exemplar B, contained 48 variants comprising 24 rotation angles under each of two mirror-flip conditions. For each stimulus set, the resulting n × 48 response matrix was divided into two n × 24 matrices and stacked along the neural-feature dimension, with corresponding rotation angles aligned, yielding a 2n × 24 pseudo-bilateral response matrix. This procedure treated responses to the two mirror-flip conditions as separate population features under an assumed mirror symmetry between hemispheres. For each anagram family, neural responses to the 72 conditions were normalized to the [0, 1] range. Spearman correlation distances between population responses at each pair of angles were used to quantify “neural” distances between conditions (Figure 3A). Two-dimensional multidimensional scaling (MDS) was applied to the RDM using MATLAB’s mdscale function. For Figures 3C and 3D, raw Spearman correlation coefficients were used for visualization, whereas Fisher z-transformed values were used for statistical comparisons. Matched and non-matched conditions were compared using t-tests across 20 interpretation pairs.

Decoding analysis: time-resolved decoding was performed at 5-ms resolution using a 50-ms sliding boxcar window spanning −50 to 400 ms relative to stimulus onset. Time-averaged decoding was computed from responses averaged over a 60–400 ms window. At each time point, population responses were z-scored to have zero mean and unit variance across trials to normalize firing rates. Trials from the two exemplar categories were pooled to form the training set and assigned binary class labels. A linear classifier was trained using L2-regularized logistic regression (fitclinear, MATLAB) to estimate the probability of category membership with lambda = 0.001. The decoder was trained on responses to all 96 variants of the canonical exemplar stimuli, so that both exemplar classes were represented across all sampled rotations and mirror configurations. The trained model was then applied to responses evoked by anagram stimuli to compute posterior probability scores for all trials. This procedure was repeated independently at each time point, yielding a time-resolved decoding trace that quantifies the probability of the population response being classified as one of the two categories for each anagram sample and time bin. Decoding scores were assessed using one- or two-sample t-tests, with P values corrected using FDR.

### Model analysis

We analyzed 37 pretrained visual models using the image preprocessing associated with each released checkpoint. For each model, we extracted representations from its final pooled, projection, or head layer. Spatial feature maps were averaged across spatial positions. The model set included 7 language-supervised models, 13 self-supervised models, and 17 image-classification models. For architecture-controlled comparisons, we further analyzed subsets of 11 Vision Transformer Base (ViT-B) models and 13 ResNet models, as shown in Figure 5.

Decoding analysis followed the same L2-regularized logistic classification procedure used for the IT data, including the same regularization parameter and decoding bias. Because model representations have no trial-by-trial variability, the decoder was trained on the 48 variants of each canonical exemplar and tested on the corresponding anagram representations. For direct comparison, neural responses were first averaged across trials and then pooled across recording sessions into a pseudo-population before applying the same decoding procedure. This averaging and pooling produced higher decoding scores than the single-trial, session-wise analysis in Figure 4. Decoding bias was averaged across the 20 family-by-canonical-angle conditions.

Shape-texture preference was measured with an independent factorial stimulus set containing 30 shapes rendered with 41 textures, for a total of 1,230 images. A shape hypothesis representational dissimilarity matrix assigned distance 0 to image pairs sharing shape and 1 to other pairs; a texture hypothesis matrix was defined analogously, as shown in Figure 5B. Model features were L2-normalized for each image, and model dissimilarities were calculated using cosine distance. We calculated the partial Spearman correlation between each model dissimilarity matrix and the shape hypothesis while controlling for texture, and the complementary partial correlation with texture while controlling for shape. Relative shape preference was defined as partial shape correlation minus partial texture correlation. Positive values indicate stronger shape-related than texture-related representation.

Single-unit tuning analysis in the models also followed the similar procedure used for the neural data, including the definition of canonical-exemplar selectivity, a selectivity threshold of 0.5, von Mises fitting, and a correlation threshold of 0.3 for retaining qualified fits. For each qualified unit, we calculated the shortest circular distance between its preferred angle and the canonical angles of the matching and non-matching exemplars. The normalized matching-angle advantage was defined as the alternative-angle distance minus the matching-angle distance, divided by the separation between the two canonical angles (90° or 180°, depending on the anagram). The resulting value ranges from -1 to 1, with positive values indicating tuning closer to the matching canonical angle and negative values indicating tuning closer to the alternative canonical angle.

**Table S1.** Summary of all recording sessions in the initial and the main experiment. “Trials” shows total trial number, with parentheses indicating average repetitions per image. “Reliable units” counts units with reliability greater than 0.4. “Initial” experiment indicates data shown in Figure S2, “Main” experiment indicates data shown in the main manuscript.

| subject | target | experiment | #image | #trials | #reliable units | #body units | #face units |
| --- | --- | --- | --- | --- | --- | --- | --- |
| F | MSB | Initial | 612 | 4313 (7.0) | 356 | 123 (0.35) | 16 (0.04) |
| Z | AL & ASB | Initial | 612 | 6648 (10.9) | 444 | 123 (0.28) | 112 (0.25) |
| J | AL & ASB | Initial | 612 | 7825 (12.8) | 530 | 154 (0.29) | 196 (0.37) |
| J | AL & ASB | Initial | 612 | 5010 (8.2) | 412 | 81 (0.20) | 118 (0.29) |
| Z | MSB | Initial | 612 | 7399 (12.1) | 231 | 55 (0.24) | 13 (0.06) |
| J | AL & ASB | Initial | 612 | 6180 (10.1) | 391 | 128 (0.33) | 96 (0.25) |
| M | ML | Initial | 612 | 4910 (8.0) | 391 | 20 (0.05) | 116 (0.30) |
| M | MF & MSB | Initial | 612 | 4966 (8.1) | 252 | 39 (0.15) | 89 (0.35) |
| M | MF & MSB | Initial | 612 | 4976 (8.1) | 318 | 43 (0.14) | 120 (0.38) |
| Z | AL & ASB | Initial | 612 | 4877 (8.0) | 299 | 43 (0.14) | 202 (0.68) |
| M | MF & MSB | Initial | 612 | 5599 (9.1) | 367 | 50 (0.14) | 115 (0.31) |
| Z | MSB | Initial | 612 | 6208 (10.1) | 459 | 231 (0.50) | 32 (0.07) |
| L | MF & MSB | Initial | 612 | 4234 (6.9) | 362 | 188 (0.52) | 40 (0.11) |
| L | MF & MSB | Initial | 612 | 4603 (7.5) | 380 | 88 (0.23) | 77 (0.20) |
| J | AL & ASB | Main | 1476 | 9858 (6.7) | 511 | 88 (0.17) | 219 (0.43) |
| J | AL & ASB | Main | 1476 | 14111 (9.6) | 259 | 22 (0.08) | 152 (0.59) |
| J | AL & ASB | Main | 1476 | 14902 (10.1) | 431 | 64 (0.15) | 192 (0.45) |
| Z | AL & ASB | Main | 1476 | 5224 (3.5) | 219 | 24 (0.11) | 155 (0.71) |
| J | AL & ASB | Main | 1476 | 9093 (6.2) | 386 | 44 (0.11) | 206 (0.53) |
| J | AL & ASB | Main | 1476 | 9049 (6.1) | 368 | 89 (0.24) | 86 (0.23) |
| C | ASB | Main | 1476 | 11976 (8.1) | 379 | 171 (0.45) | 10 (0.03) |
| C | ASB | Main | 1476 | 14881 (10.1) | 381 | 118 (0.31) | 17 (0.04) |
| J | AL | Main | 756 | 8829 (11.7) | 284 | 3 (0.01) | 202 (0.71) |
| J | AL | Main | 756 | 8785 (11.6) | 302 | 7 (0.02) | 192 (0.64) |
| J | AL & ASB | Main | 756 | 7789 (10.3) | 536 | 58 (0.11) | 253 (0.47) |
| Z | AL & ASB | Main | 756 | 7608 (10.1) | 432 | 124 (0.29) | 168 (0.39) |
| J | AL | Main | 756 | 8651 (11.4) | 237 | 6 (0.03) | 176 (0.74) |

**Supplementary Figure S1:**
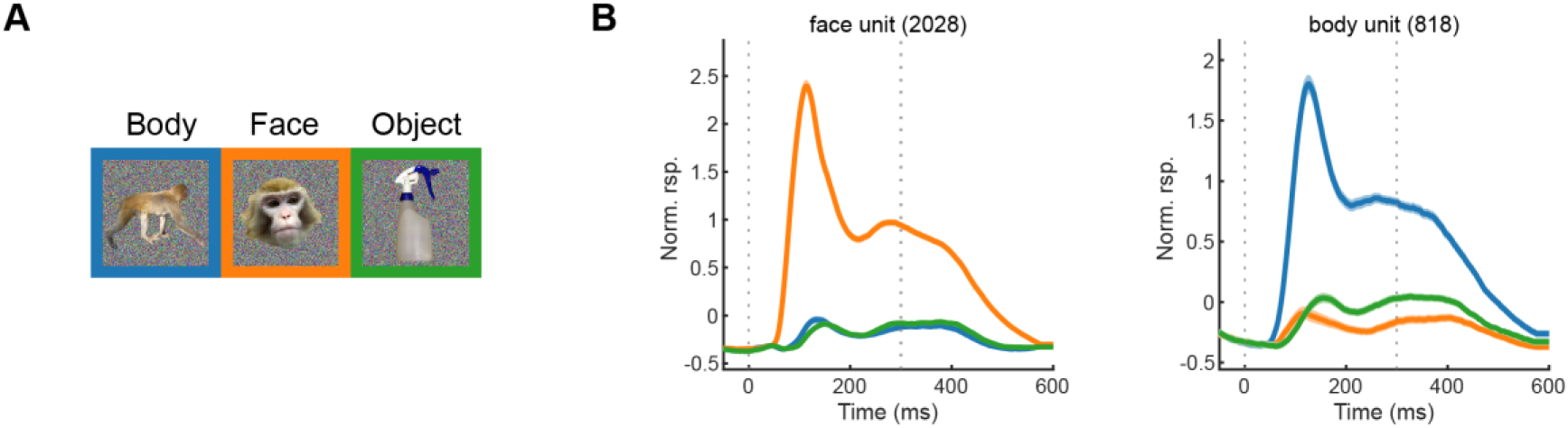
Summary of recorded face- and body-selective units. (A) Example localizer stimuli. (B) Population-averaged, normalized PSTHs of face-selective and body-selective units in response to localizer stimuli. Shaded areas indicate 95% confidence intervals for the mean across units. Vertical dashed lines denote stimulus onset and offset. Note that although we collected data from face- and body-selective areas, the main analyses used selectivity defined by the contrast between the two exemplars.

**Supplementary Figure S2:**
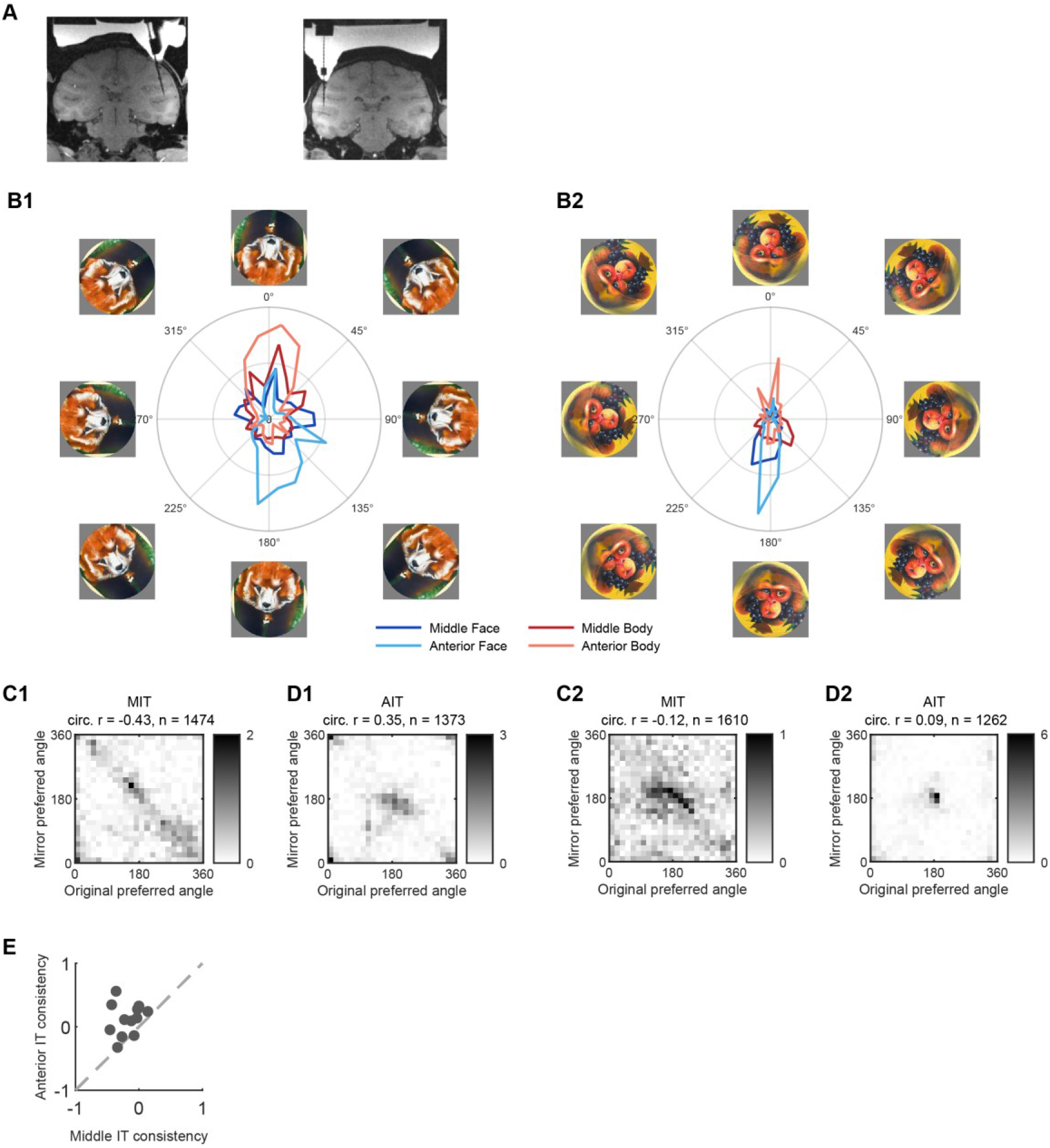
Results from an initial experiment showing greater preferred-angle consistency in anterior than middle inferotemporal cortex. The initial experiment comprised 14 recording sessions from five macaques. Nine sessions targeted middle IT (MIT) and yielded 3,116 reliable neural units, five sessions targeted anterior IT (AIT) and yielded 2,076 reliable units. In total, 5,192 units passed the initial reliability criterion (Spearman–Brown-corrected split-half reliability > 0.4). (A) Representative middle IT recording sites from two additional monkeys, F and L. (B) Representative stimuli from two anagram families at eight angles, along with polar distributions of preferred angles in face- and body-selective units in middle and anterior IT. Blue denotes face-selective units and red denotes body-selective units; darker and lighter shades denote middle and anterior IT, respectively. (C, D) Binned-density distributions of preferred-angle estimates in MIT (C) and AIT (D). The horizontal axis shows the preferred angles fitted for the original condition, and the vertical axis shows the preferred angles fitted for the mirrored condition. Each bin indicates the percentage of units. For both example families, horizontal mirroring produced less consistent preferred angles in MIT than in AIT, as reflected by lower circular–circular correlations between the two estimates in MIT. (E) Comparison of circular–circular correlations between MIT and AIT across the 12 anagram families. Each point represents one anagram family, the dashed diagonal denotes equal consistency in the two regions. Preferred-angle consistency was greater in AIT than in MIT across families (Wilcoxon signed-rank test, P < .001, n = 12 families).

**Supplementary Figure S3:**
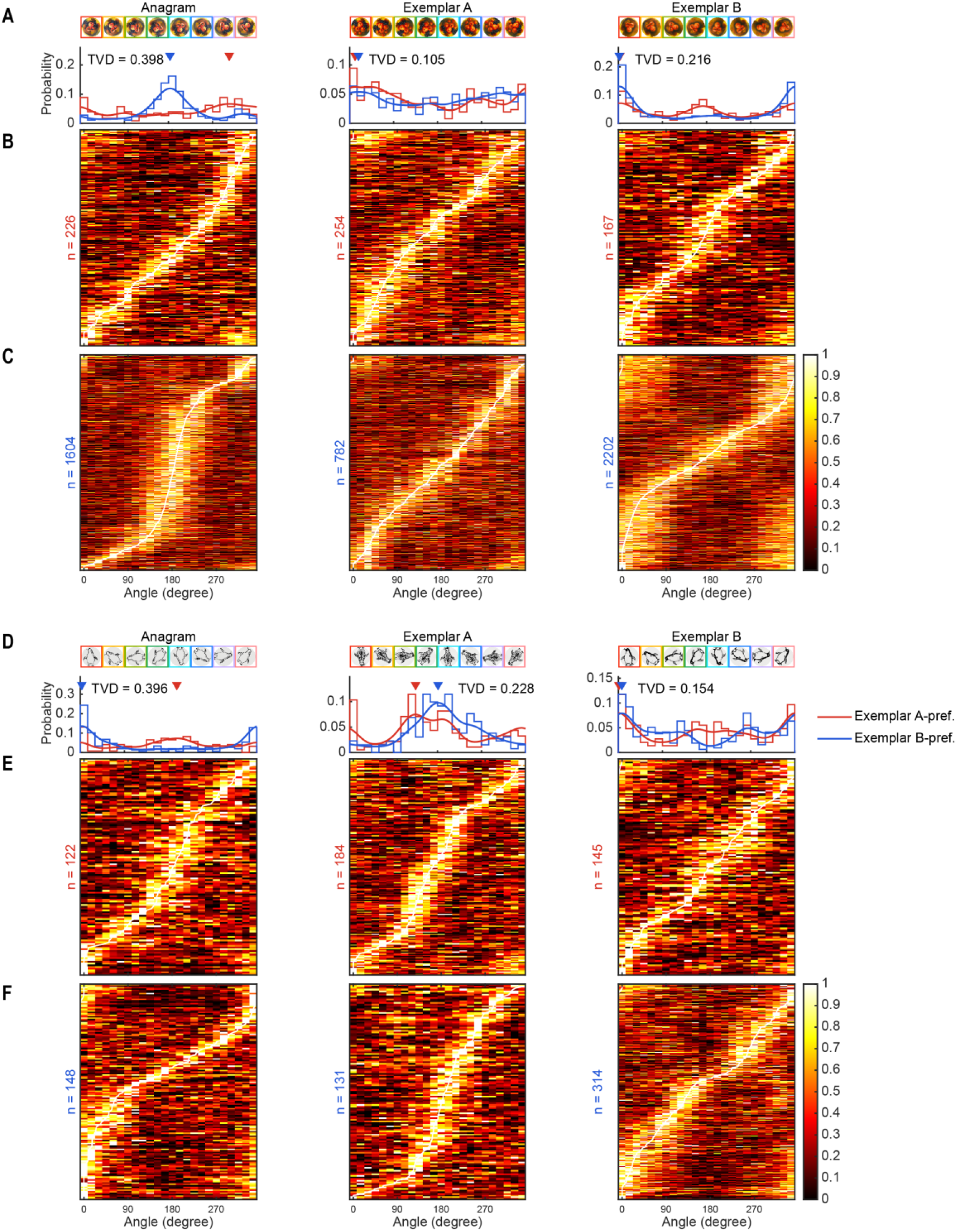
More single-unit tuning examples, related to Figure 2. (A–C) Tuning summary for the monkey–fruit anagram. (D–F) Corresponding results for the penguin–giraffe anagram. Conventions are the same as in Figure 2. (A, D) Example images and distributions of fitted preferred angles. (B, E) Heatmaps showing rotation-dependent responses of fruit-preferring and giraffe-preferring neurons, respectively. (C, F) Same as (B, E), but for monkey-preferring and penguin-preferring neurons, respectively.

**Supplementary Figure S4:**
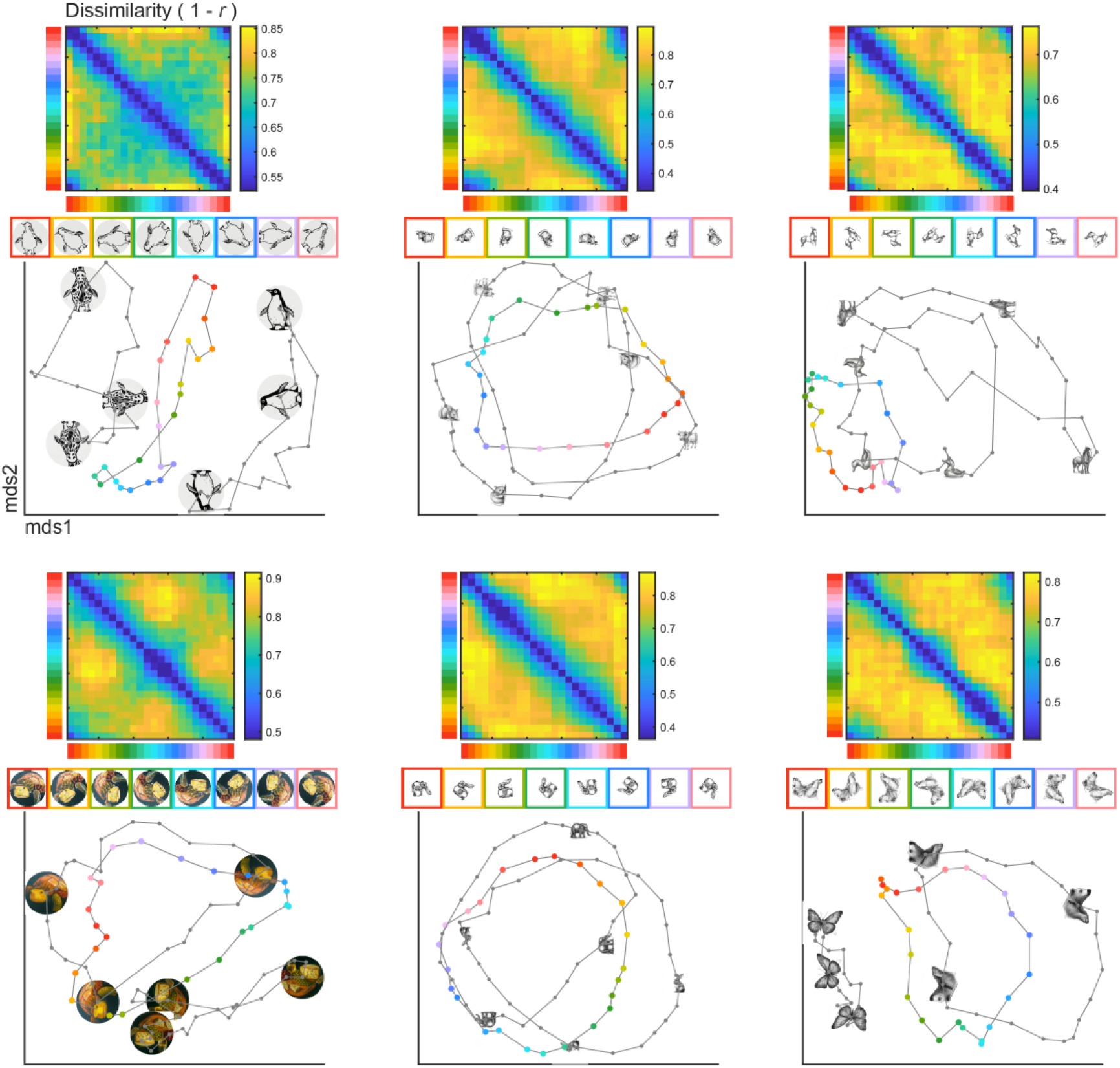
More population geometry examples, related to Figure 3. Population RDM, MDS visualization for additional anagram examples. Conventions are the same as in Figure 3.

**Supplementary Figure S5:**
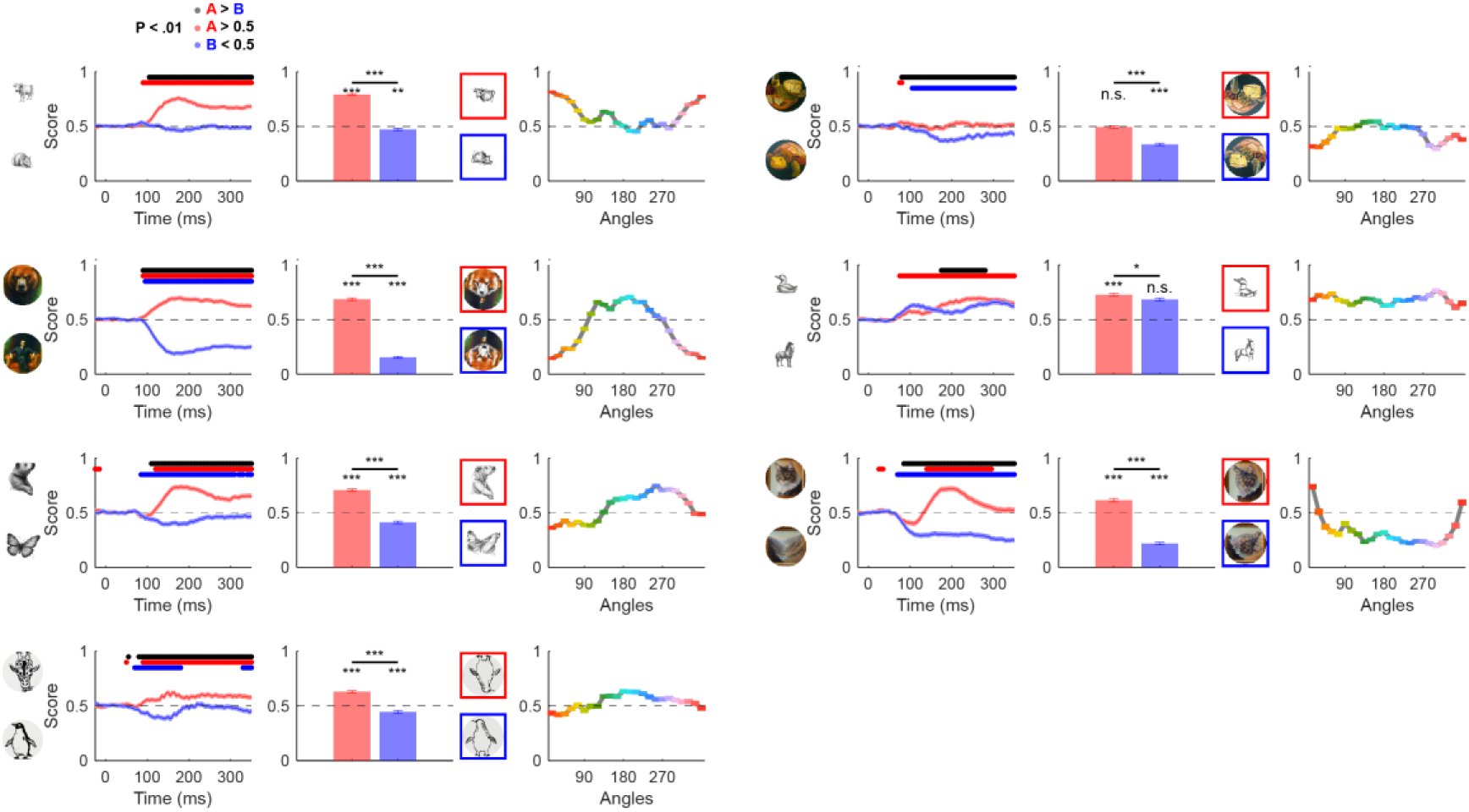
More population decoding examples, related to Figure 4. Decoding results for the additional anagram pairs. Conventions are the same as in Figure 4.

**Supplementary Figure S6:**
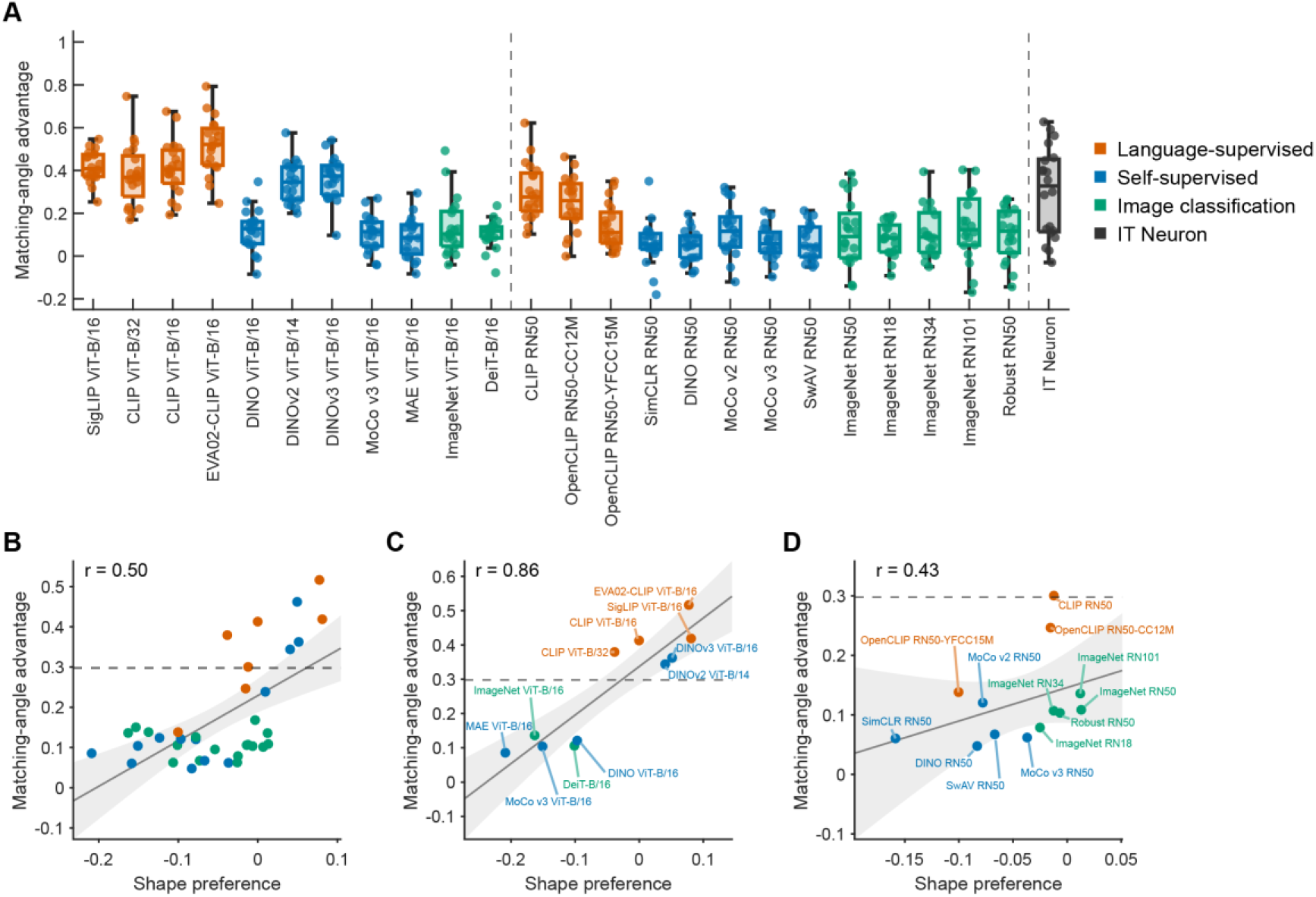
Single-unit tuning in visual models. (A) Normalized matching-angle advantage for 11 ViT-B models, 13 ResNet models, and the IT neuronal population. For each qualified tuning curve, the distance between the preferred anagram rotation and the canonical angle of the unit’s preferred canonical exemplar was compared with the distance to the alternative canonical angle. Each point represents one of the 20 interpretations. (B) Relationship between relative shape preference and normalized matching-angle advantage across all 37 models. Orange denotes language-supervised models, blue denotes self-supervised models, and green denotes image-classification models. (C) Same relationship for the 11 ViT-B models. (D) Same relationship for the 13 ResNet models.

**Supplementary Figure S7:**
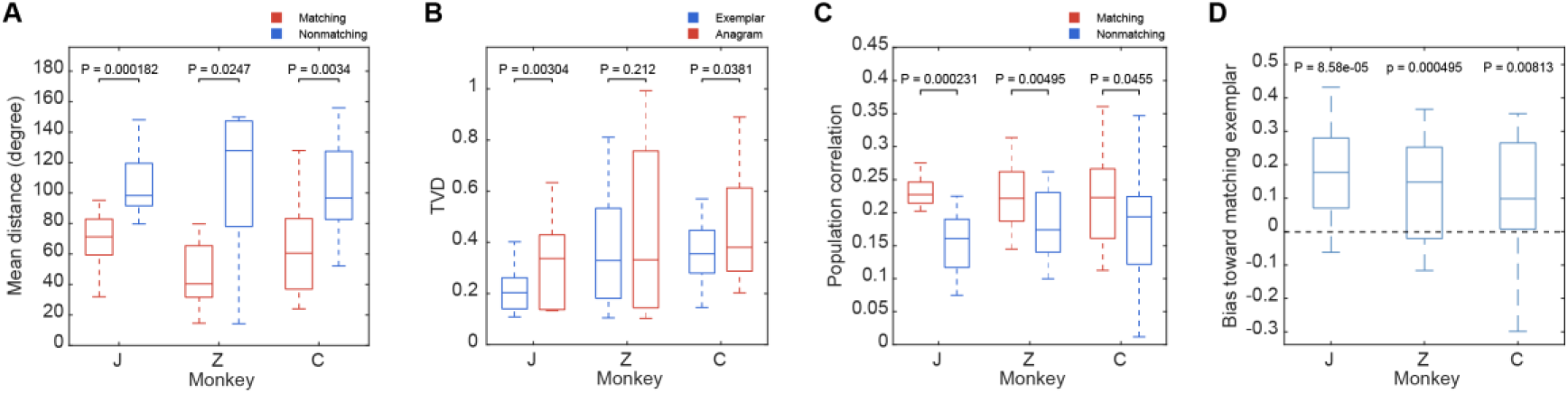
Main effects in individual monkeys, J, Z, C. (A) Related to Figure 2G, circular distance between the preferred angle of each qualifying fit and the critical angle associated with either the matching or nonmatching interpretation. (B) Related to Figure 2H, total-variation distance between the two canonical-exemplar-selective neural populations for canonical exemplars and anagrams. (C) Related to Figure 3C, mean population correlation between anagrams at interpretation-critical angles and matching or nonmatching canonical exemplars. (D) Related to Figure 4F, decoder bias toward the matching canonical exemplar.

